# A Causal and Dissociable Role for the Right Inferior Prefrontal Cortex in Empathy for Physical and Social Pain

**DOI:** 10.1101/2025.01.19.633761

**Authors:** M. De Lillo, A. Korpal, H. Ferguson, A. K. Martin

## Abstract

The right inferior frontal gyrus (rIFG) and dorsomedial prefrontal cortex (dmPFC) are key nodes in the social brain, implicated in empathy for physical and social pain. However, their causal and dissociable contributions remain unclear. In this study, 52 young adults underwent focal transcranial direct current stimulation (f-tDCS) targeting the rIFG or dmPFC in a sham-controlled, double-blind, crossover design. Participants rated the intensity of pain in images depicting social or physical pain during stimulation. Anodal stimulation to the rIFG increased ratings of physical pain and decreased ratings of social pain, suggesting dissociable roles in processing empathy for these two pain types. In contrast, dmPFC stimulation did not modulate ratings, potentially reflecting its role in higher-order social cognitive processes rather than affective empathy. The effects of rIFG stimulation on social pain were significantly stronger in the initial trials, suggesting potential habituation within the rIFG or stimulation-specific effects. These results provide causal, dissociable evidence for the rIFG’s involvement in empathy, with its effects differing based on the type of pain. This supports the proposal that distinct neural processes underlie empathy for social versus physical pain.

---

Empathy refers to the ability to understand and share the mental states of others (Singer et al., 2004) and is the process through which we comprehend both the emotional and physical states of others, including their experience of pain (Decety & Jackson, 2004; Ferguson & Wimmer, 2023; Gallese, 2003; Timmers et al., 2018). Empathy can often be felt for others in physical pain, but we also empathise with people experiencing social pain, such as bereavement or social rejection. Considerable research has explored the shared and unique neural mechanisms underpinning processing of physical and social pain in relation to oneself (Eisenberger et al., 2012a; Meyer et al., 2015; Ochsner et al., 2008). However, much less is known about the extent to which common and unique neural processes are involved in understanding physical and social pain experienced by others. Causal evidence for the role of key social brain regions in relation to processing physical and social pain in others is lacking. In this paper, we address this gap directly by using focal transcranial direct current stimulation to excite either the right inferior frontal gyrus (rIFG) or the dorsomedial prefrontal cortex (dmPFC) whilst participants rated the physical or social pain experienced by others.

Much of our understanding of the empathic brain has been acquired through neuroimaging studies that have focussed on empathy for physical pain (Masten et al., 2011; Singer et al. 2004). This work has shown that empathizing with others experiencing physical pain recruits brain regions such as the anterior insula (AI), the dorsal anterior cingulate cortex (dACC) and neighbouring cortical regions, such as the right inferior frontal gyrus (rIFG; Lamm et al., 2010; Masten et al., 2011; Singer et al., 2004). The rIFG is a region that is consistently associated with processing incoming bodily signals. It is involved in processing both pain to oneself and with empathic responses to physical pain in others (Budell et al., 2010, 2015; Iacoboni, 2009; Li et al., 2021; Vachon-Presseau et al., 2012). The IFG is also considered a key hub of the human mirror neuron system (hMNS) and as such, has been associated with the affective component of empathy (Shamay-Tsoory et al., 2009; Wu et al., 2018). In sum, the IFG is a brain region that is crucial for understanding others’ actions, intentions, thoughts and emotions.

Research into the empathic response to social pain in others has received less attention. Social pain includes painful experiences, such as social exclusion or rejection (Williams. 2007). Some researchers have conceptualised social pain as an emotional response to recognising a psychological distance from an individual’s desired social connections (Eisenberger, 2012b; Eisenberger, 2015). Others propose that it reflects the distress caused when an individual’s primary human need to feel a sense of belonging, control and meaning goes unmet (Williams, 2007). Very little is known about how this is processed in the brain. One potential brain is the dorsal medial pre-frontal cortex (dmPFC), which is a vital hub in the social brain (Schurz et al., 2014) implicated in a diverse range of social cognitive processes, including Theory of Mind (ToM; Spunt & Adolphs, 2017), perspective-taking (Martin et al., 2017), and self-other processing (Martin et al., 2019; Wittmann et al., 2016). The dmPFC may also have a crucial role in interpreting the cognitive component of empathy (Masten et al., 2011) as well as empathy for the social suffering of strangers (Meyer et al., 2013).

An unresolved question among empathy researchers is the extent to which neural processes are shared when processing physical and social pain in others (Eisenberg, 2015; Ferguson et al., 2024; Flasbeck & Brüne, 2019; Iannetti et al., 2013). Some studies have shown that overlapping neural areas are recruited in response to others in social and physical pain, despite their diverging phenomenology (Ferguson et al., 2024). Other studies support dissociable roles for these brain regions, with the dorsomedial prefrontal cortex (dmPFC) potentially involved in processing social pain to a greater extent than physical pain (Bruneau et al., 2013; Meyer et al., 2015). The physical-social pain overlap hypothesis (Panksepp, 1998) suggests that social pain relies on some of the same neural regions involved in processing physical pain, particularly the anterior insula (AI) and dorsal anterior cingulate cortex (dACC), as well as neighbouring cortical regions such as the right inferior frontal gyrus (rIFG) and the dmPFC (Masten et al., 2011). However, other researchers have contested the notion that these systems are completely homogenous (e.g., Eisenberger, 2015). Shared neural networks for processing social and physical pain may have evolved as an adaptive mechanism, promoting behaviours that help individuals respond effectively to threats or challenges (Sturgeon & Zautra, 2016). Despite this, most research has focused on self-experienced social and physical pain, with less attention given to how these networks contribute to empathizing with others’ pain.

Recent studies have begun addressing this gap by examining empathy for physical and social pain within matched experimental paradigms. Ferguson et al. (2024) used EEG and a pain-rating task with carefully matched stimuli to investigate neural responses associated with empathy for physical and social pain. They found greater mu desynchronization, an EEG marker linked to the human mirror neuron system (hMNS), in response to pain versus no-pain situations, but no significant differences between physical and social pain. In contrast, Flasbeck and Brüne (2019) identified distinct early (330–450 ms) and late (500–700 ms) event-related potential (ERP) responses, with physically painful stimuli eliciting stronger late ERP responses than socially painful stimuli, particularly in components such as the late positive potential (LPP). These studies highlight the temporal dynamics of empathic processes but reveal limitations in determining causal relationships between specific brain regions and empathy. Recent advances in non-invasive brain stimulation have provided causal evidence for the role of the rIFG and related regions in empathy. For instance, Wu et al. (2018) showed that excitatory stimulation to the rIFG increased cognitive empathy, while He et al. (2020) demonstrated that stimulating the rVLPFC reduced negative emotional reactions to social pain. However, neither of these studies tested empathy for both physical and social pain within the same task and cohort, and methodological limitations such as non-targeted stimulation and lack of control sites, constrains the generalizability of findings.

The present study aims to provide causal evidence for the role of the right inferior frontal gyrus (rIFG) and the dorsal medial prefrontal cortex (dmPFC) in empathy for social and physical pain in others. These regions were selected as they represent key hubs in the social brain network, with potentially complementary roles in processing the cognitive and affective dimensions of empathy (Bruneau et al., 2013; Lamm et al., 2007; Masten et al., 2011; Meyer et al., 2015; Singer et al., 2004). By employing focal transcranial direct current stimulation (f-tDCS), this study overcomes the limitations of diffuse stimulation methods, allowing us to isolate the contributions of these specific regions to physical and social empathy. This investigation directly addresses a key unresolved question in empathy research: to what extent do neural processes overlap when processing physical and social pain in others? We hypothesize that stimulation of the rIFG will enhance empathic responses to physical pain, given its established role in processing bodily signals and affective empathy, and may simultaneously reduce ratings of social pain, potentially reflecting its involvement in down-regulating emotional responses to social distress. Conversely, we predict that stimulation of the dmPFC will preferentially modulate empathic responses to social pain, aligning with its involvement in interpreting complex social interactions and mental states.

This study will provide critical insights into the functional specialization and interaction of these regions within the broader empathy network. By testing whether stimulation of these regions produces overlapping or dissociable effects on physical and social pain empathy, the study informs ongoing debates about the shared versus unique neural mechanisms that underlie these processes. If overlapping effects are observed, this would support the physical-social pain overlap hypothesis, suggesting evolutionary and functional commonalities. Alternatively, dissociable effects would indicate distinct neural pathways for processing physical and social pain, reinforcing the idea that these experiences engage unique cognitive and affective mechanisms. By isolating the contributions of these brain areas, this research advances our understanding of empathy’s neural architecture and offers potential avenues for future targeted interventions in clinical contexts involving impaired social or emotional processing.

## Methods

### Participants

Power calculations showed that 52 participants were required (26 in each stimulation condition) for the present study to have 80% power to detect a large effect (cohen’s f = 0.4), with a significance level of .05, comparable to previous research using f-tDCS to modulate social cognition (Martin, Huang, et al., 2019). The participants were 52 healthy young adults (aged between 18-35 years old), stratified to receive stimulation over either the dmPFC (n = 26) or rIFG (n = 26) in sham-controlled, double-blinded, crossover studies. Stimulation order was balanced across both stimulation sites so that half of the participants received active and half received sham stimulation during their first session. The groups were comparable in age, gender, autism-quotient scores (Baron-Cohen et al., 2001), and hospital depression and anxiety scores (Zigmond & Snaith, 1983; see Table 1). All participants were tDCS naïve, were not currently taking psychoactive medication or substances, and had no history of neurological or severe psychiatric issues. All participants provided written consent, completed a safety screening questionnaire before testing, and were compensated for their time with course credits or a small monetary compensation. The study abided by the ethical standards as per the 1991 Declaration of Helsinki and ethical approval was granted by the Research Ethics Committee within The School of Psychology at The University of Kent.

**Table 1.** Demographics across stimulation condition groups. P-value shows between-group comparisons.

|  | rIFG | dmPFC | *p* |
| --- | --- | --- | --- |
| Age (years) | 22.16 (3.60) | 21.42 (3.69) | .50 |
| Gender | 20F/6M | 19F/7M | .75 |
| Autism Quotient | 20.85 (8.65) | 19.81 (9.94) | .69 |
| HADS-depression | 5.58 (3.67) | 6.46 (3.85) | .40 |
| HADS-anxiety | 7.81 (4.75) | 8.00 (4.49) | .88 |

#### Pain Rating Task

We used a modified version of the pain rating task used in Ferguson et al. (2024), which was based on the task used in Jackson et al. (2005), presented using PsychoPy version 2022.1.3. The task involved the presentation of 80 images (taken directly from Ferguson et al., 2024): 20 images depicted no physical pain, 20 images depicted physical pain, 20 images depicted no social pain and 20 images depicted social pain (see https://osf.io/f9c2r/ for full set of images). Participants completed four blocks resulting in a total of 320 trials (each image was presented four times in total), with rest breaks provided between each block.

Each trial began with a fixation cross for 500ms followed by an image for 3,500ms. After the presentation of each image, a screen prompted participants to rate “How painful was this situation?” (i.e. the level of pain the person in the picture was feeling). Responses were made on a visual analogue scale from 0 (no pain) to 100 (worst pain) using a mouse. A blank screen was presented for 500ms between trials. Social and physical pictures were presented in separate blocks, with the order counterbalanced (for each cohort based on brain region) so that half received social images first and the other half received physical images first (see Figure 1 for task design and timings). The pain rating task took approximately 30 minutes to complete in each session.

**Figure 1.**
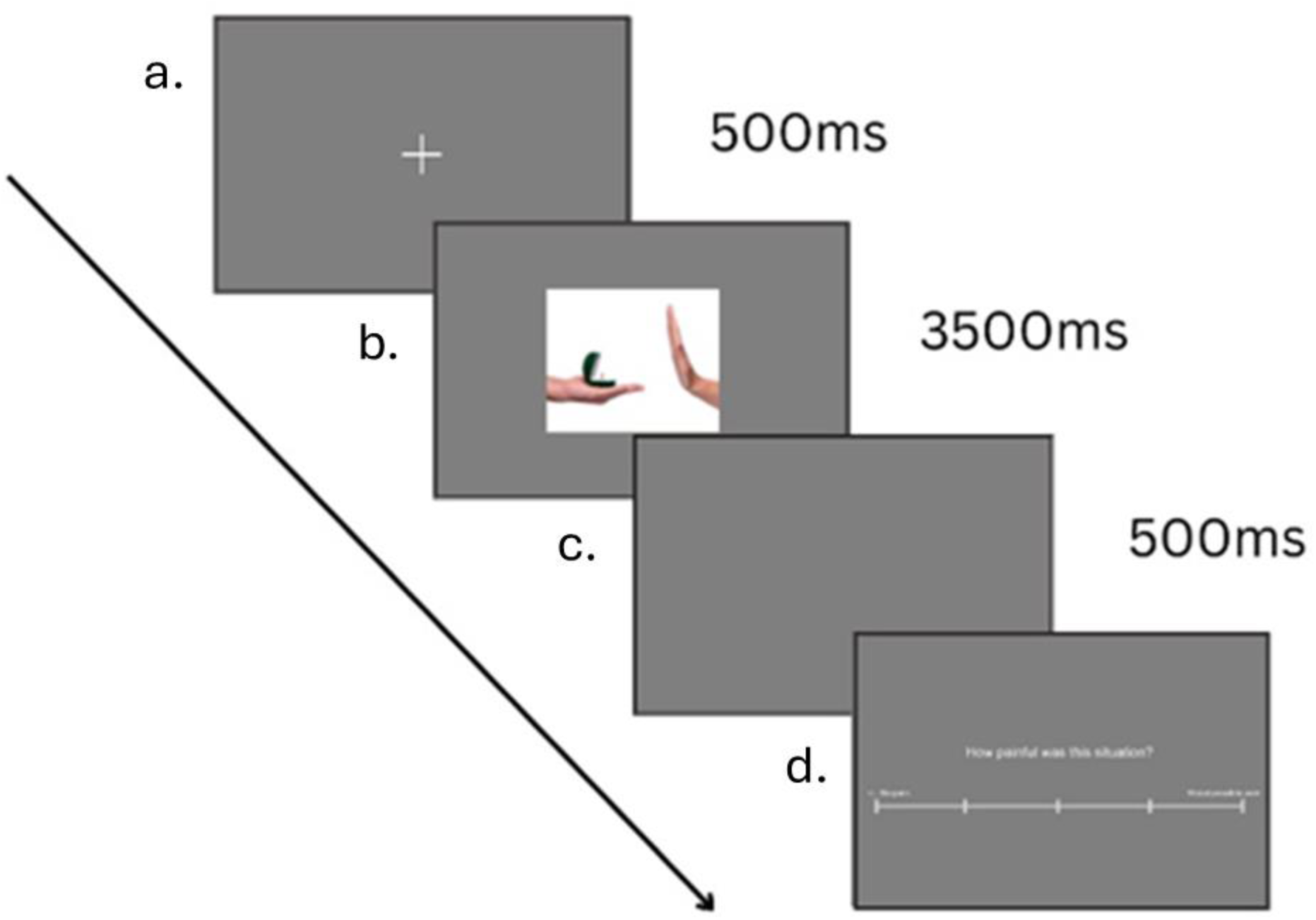
Schematic illustration of the task design and timings. Note. a. = fixation cross (500ms), b. = stimulus (3,500ms), c. = blank screen (500ms), d. = rating (“How painful was the situation?”).

### Focal-tDCS

Focal-tDCS was administered to participants using a one-channel direct current stimulator (DC-Stimulator Plus, NeuroConn) and two concentric rubber electrodes (Bortoletto et al., 2016; Gbadeyan, McMahon, et al., 2016). The small centre electrode was 2.5cm in diameter, and the ring-shaped return electrode had a diameter of 7.5/9.8 cm (inner/outer). The setup is similar to the ‘4 x 1’ focal-tDCS setup in which the current is constrained by utilising four return electrodes arranged in a circle around the centre electrode (Alam et al., 2016; Hogeveen et al., 2016; Kuo et al., 2013). The present setup was preferred as it does not require an expensive multi-channel simulator and has been shown to provide similar focal current delivery (Martin, Huang, et al., 2017, 2019; Martin, Su, et al., 2019). Secure electrode placement over target regions was achieved using an adhesive conductive gel (Weaver Ten20VR conductive paste) and electrodes were held in place using an EEG cap to ensure stable conductive adhesion to the skin. The location of the centre electrode when stimulating the dmPFC and rIFG was determined using the 10–20 international EEG system. The scalp region overlying the dmPFC was located by measuring 65% of the distance from the Cz and the Fpz (Martin, Dzafic, et al., 2017). The central electrode was placed at FC6 for rIFG stimulation (Hogeveen et al., 2015, 2016). The ring electrode was placed equidistantly around the centre electrode at each brain region.

During the active stimulation condition, the current was ramped up to 1 mA over 5 seconds, with the focal-tDCS being administered for 20 minutes before ramping down over 5 seconds. In the sham stimulation condition, there was a ramp-up period of five seconds, but stimulation was only administered for forty seconds before ramping down over a period of five seconds. The sham condition mimics the physical sensation of active stimulation and has been shown to effectively blind participants as to which stimulation they received (Gbadeyan, Steinhauser, et al., 2016; Martin, Dzafic, et al., 2017; Martin, Huang, et al., 2019; Martin, Su, et al., 2019) with no neurophysiological effect (Stagg et al., 2013). The study was double-blinded as the researcher was blinded to the experimental condition by utilising the ‘study mode’ of the DC-stimulator (i.e., a pre-assigned code triggered the respective stimulation conditions). All impedances were below 55 kΩ at the beginning of stimulation as per safety requirements. Stimulation was counterbalanced so that half of participants received sham stimulation first and the other half received active stimulation first.

Current modelling was conducted using SimNIBS version 4.1 (Thielscher et al., 2015; see Figure 2). Simulation parameters were chosen to reflect the experimental setup of this study (i.e., intended positions of the centre anode and ring cathode, electrode dimensions, current strength, gel thickness). Electrode thickness and gel thickness were defined as 2mm and 1mm, respectively. Standard conductivity values as provided by SimNIBS were used. For the dmPFC montage, the centre position of the anode was initially determined in MNI space by identifying FPz and Fz and measuring the distance between the two points. The target scalp coordinates were then determined by measuring 15% of the distance from Fz towards FPz (MNI scalp coordinates: 0.5/71.7/46.1). The center position of anode for the rIFG montage corresponded to the FC6 position of the EEG-10-10 system provided by SimNIBS. Ring cathodes were placed equidistantly around the central anode. Figure 2 shows the theoretical cortical electric field (magnitude E) for anodal focal-tDCS to both the rIFG and the dmPFC. Peak E-Field was 0.13 v/m for rIFG and 0.09 v/m for dmPFC.

**Figure 2.**
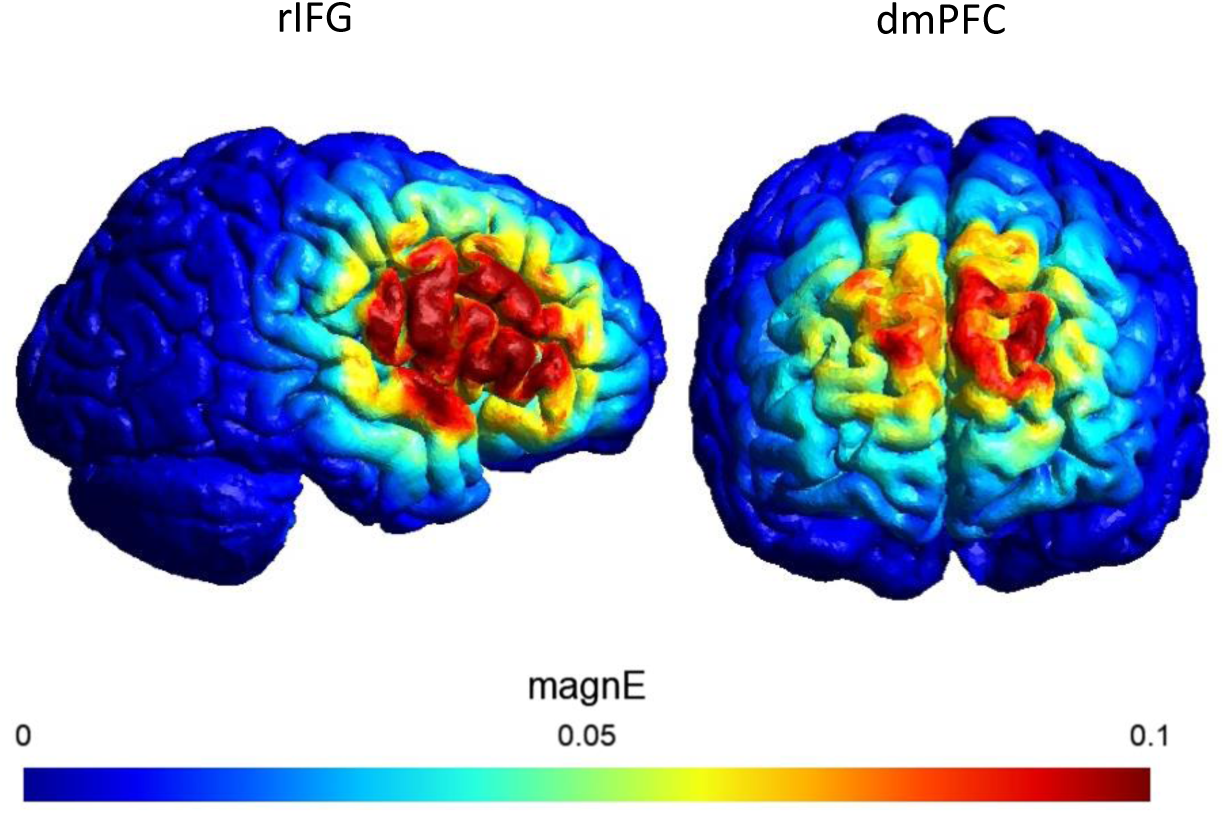
Theoretical electric field to the dmPFC and rIFG during online anodal focal-tDCS.

#### Questionnaires

##### Autism Quotient

The Autism-Spectrum Quotient (AQ; Baron-Cohen et al., 2001) is a self-administered questionnaire consisting of 50 items designed to measure the degree to which adults exhibit traits associated with the autism spectrum. It returns a score between 0-50 with higher scores indicating a greater presence of autism-related traits.

##### Hospital Anxiety and Depression Scale

The Hospital Anxiety and Depression Scale (HADS; Zigmond & Snaith, 1983) is a self-administered questionnaire to assess anxiety and depression levels. It comprises 14 items, with seven items each dedicated to anxiety (HADS-A) and depression (HADS-D). Each item is scored on a 4-point Likert scale (0–3), resulting in subscale scores ranging from 0 to 21. Higher scores indicate greater levels of anxiety or depression.

#### Mood change and adverse effects

Positive and negative mood was assessed before and after each stimulation session using the visual analogue of mood scales (VAMS, Folstein & Luria, 1973). Participants were presented with a series of mood descriptors, (happy, energetic, afraid, tense, angry, tired, confused, sad) and were asked to indicate their mood intensity for each descriptor by placing a mark on a horizontal line. The line was presented between a neutral emoji and the emoji representing the emotions listed above. The scores were calculated based on the distance of the participant’s mark from the starting point of the line.

Adverse effects were assessed following each stimulation session (Brunoni et al., 2011). Participants rated the following side effects of stimulation using a scale between 1 (absent) and 4 (severe): headache, neck pain, scalp pain, tingling, itching, burning sensation, skin redness, sleepiness, trouble concentrating, acute mood change, other. We summed across these to compute a total adverse effects score.

#### Procedure

Participants completed two lab-based testing sessions, a minimum of three days apart. During the first session, participants were given full information about the project and provided informed consent. Next, they completed a demographics and safety screening form. Once eligibility and safety were assured, participants completed the AQ, HADS and VAMS questionnaires. TDCS commenced at the onset of the pain rating task. After pain rating task, participants completed the VAMS again and a questionnaire about adverse effects. In session two, participants completed the VAMS questionnaire before and after the pain rating task, and the adverse effects questionnaire after the pain rating task. Upon completion of the second session, to assess whether successful blinding was achieved, participants were asked to indicate which session they believed was the real stimulation. Finally, participants were debriefed and compensated for their time.

#### Statistical Analysis

All analyses were computed in JASP version 0.18.3 (JASP Team, 2024) or R (R Core Team, 2023; version 4.3.2). To assess the effects of stimulation to either the rIFG or the dmPFC on social and physical pain ratings, we computed a 2×2×2×2 mixed analysis of variance (ANOVA) with pain rating (0-100) as the outcome variable, Brain Region (rIFG vs dmPFC) as a between-subjects factor, and Stimulation (sham vs anodal), Type (pain vs no-pain) and Content (social vs physical) as within-subjects factors.

## Results

Blinding was achieved as participants were unable to guess the real stimulation session better than chance at either the rIFG (50% guessed correctly, χ²(1) = 0.00, *p* = 1.0) or the dmPFC (58% guessed correctly, χ²(1) = 0.62, *p* = .43).

### Pain Ratings

Figure 3 shows pain ratings in each condition. A mixed ANOVA was conducted to assess how anodal stimulation to either the dmPFC or rIFG modulated ratings of social or physical pain. A main effect of TYPE demonstrated that participants distinguished painful from non-painful stimuli (pain > no pain), F(1, 50) = 1086.24, *p* < .001, ηₚ² = 0.96. A Type × Content interaction, F(1, 50) = 45.85, *p* < .001, ηₚ² = 0.48, showed that this pain difference was larger for physical content (*t* = 31.64, *p* < .001, *d* = 4.49) than for social content (*t* = 24.57, *p* < .001, *d* = 3.49). This interaction was further subsumed under a four-way interaction between Brain Region × Stimulation × Type × Content, F(1, 50) = 4.34, *p* = .04, ηₚ² = 0.08. To investigate this interaction, separate repeated-measures ANOVAs were conducted for each brain region.

**Figure 3.**
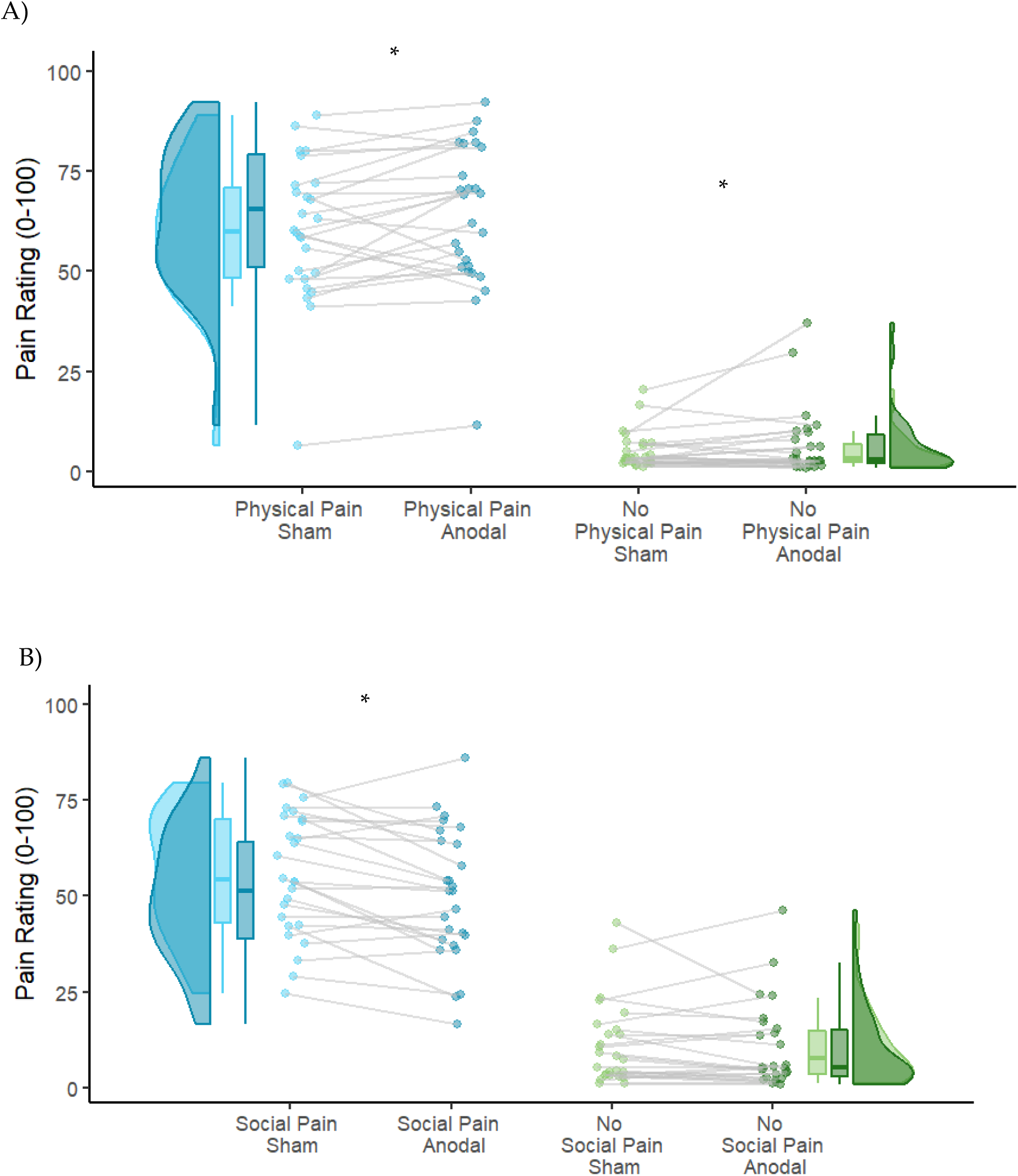
Effects of anodal stimulation to the rIFG on physical and social pain ratings. A) Stimulation increased ratings of pain in images depicting physical pain and control images depicting no physical pain. B) Anodal stimulation to the rIFG decreased rating for social pain only in images depicting social pain. * denote significant differences (p<.05).

At the rIFG, evidence supported an interaction between Stimulation x Content x Type, F(1, 25) = .6.91, *p* = .02, η_p_^2^ = 0.22. Separate analyses were conducted for social and physical content. For social content, an interaction between Stimulation and Type was identified, F(1,25)= 7.21, p=.01, η_p_^2^ =0.22. Simple effects analyses showed that anodal stimulation to the rIFG reduced pain ratings for the images depicting social pain, t=3.80, p<0.001, d=0.39 and no effect for the images depicting social no-pain, t=0.44, p=.67, d=0.05. For physical pain, The interaction between Stimulation and Type was not supported, F(1,25)= 0.60, p=.45, η_p_^2^=0.02. However it should be noted that in the overall model a Stimulation x Content interaction was also supported, F(1,50)=10.17, p=.002, η_p_^2^=0.17. In the physical pain condition, Stimulation increased pain ratings in general, F(1,25)=6.91, p=.01, η_p_^2^=0.22. Figure 5 reports pain ratings separately for the physical and social pain following rIFG stimulation.

For the dmPFC, the evidence supported the null hypothesis for all effects of stimulation. Specifically, the interaction between Stimulation x Content x Type was not supported, F(1,51)= 0.01, p=.92, η_p_^2^< 0.001. Likewise, Stimulation x Content, F(1,51)= 0.04, p=.84, η_p_^2^= 0.002, Stimulation x Type, F(1,51)= 0.08, p=.78, η_p_^2^=0.003, and the main effect of Stimulation, F(1,51)= 0.003, p=.96, n2p<0.001, were all non-significant.

Therefore, stimulation to the rIFG has dissociable effects on images depicting social and physical pain with a reduction in pain rating in images depicting social pain and a general increase in the rating of physical pain regardless of whether the image depicted pain or not.

To further explore the significant rIFG stimulation effects in opposing directions for social and physical pain, we examined how these stimulation effects unfolded over trials. We computed a linear mixed effects model using the LMER package (Bates et al., 2015) in R. P-values were obtained using the Satterthwaite approximation of degrees of freedom via the lmertest package (Kuznetsova et al., 2017). The analysis was performed separately for physical and social pain. To examine whether the relationship between Trial and Rating was better characterized by a linear or quadratic function, we compared two mixed-effects models. Both models included Stimulation as a fixed effect and random intercepts for participants. The second model additionally included a quadratic term for Trial.

For social content, model comparison using a likelihood ratio test revealed that the quadratic model provided a significantly better fit to the data (χ²(1) = 11.72, *p* < .001). The Akaike’s Information Criterion (AIC) values also supported the quadratic model (AIC = 38869) over the linear model (AIC = 38879). These results suggest that the relationship between Trial and Rating follows a quadratic trajectory rather than a linear one. For social content, the Stimulation x Trial interaction was significant, ß = -3.87, SE = 0.79, t(1, 4130.2) = -4.90, *p* < .001, showing that stimulation had a disproportionately stronger effect on reducing social pain ratings among earlier trials. The difference in social pain ratings during sham and anodal sessions, reduced by 3.87 with each subsequent trial.

For physical content, model comparison using a likelihood ratio test revealed that the quadratic model did not provide a significantly better fit to the data (χ²(1) = 0.07, *p* = .79). The AIC values also supported the linear model (AIC = 37578) over the quadratic model (AIC = 37580). These results suggest that the relationship between Trial and Rating is best characterized as linear rather than quadratic. For physical content, the Trial x Stimulation interaction was not significant, ß = 0.01, SE = 0.01, t(1,4131.1) = 1.10, *p* = .28, but the effect of Trial was significant, ß = -0.02, SE = 0.01, F(1,3973.7) = -2.91, *p* = .004, showing that pain ratings increased in parallel over trials for anodal and sham stimulation. Pain ratings over trials are plotted for each stimulation condition and social/physical content in Figure 4.

**Figure 4.**
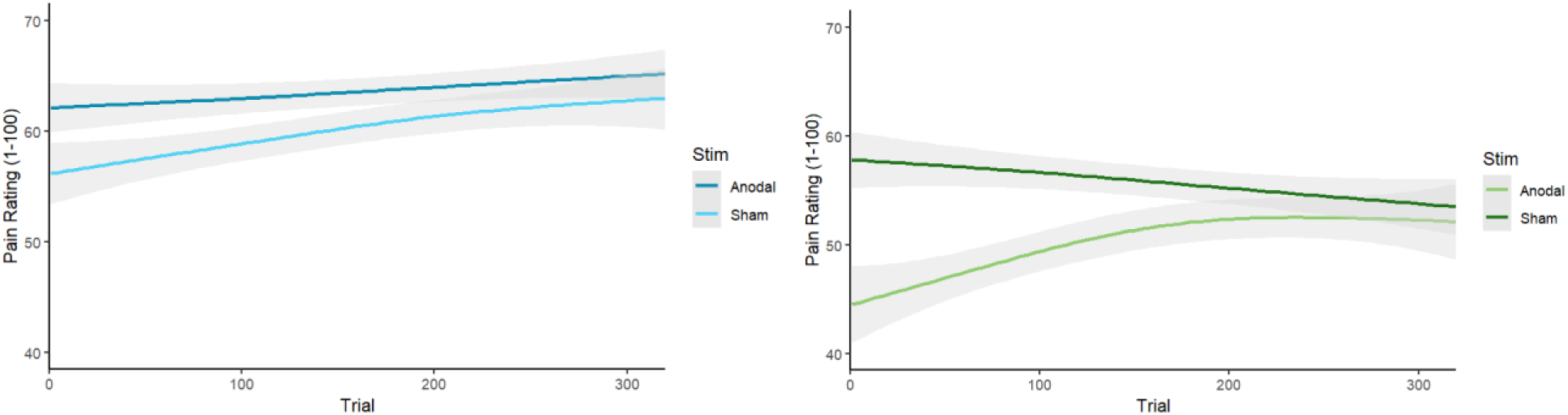
Effect of anodal stimulation to the rIFG on physical (left panel) and social (right panel) pain over trials. **Note.** Anodal stimulation to the right IFG reduced social pain rating disproportionately in the earliest presentations of painful images. In contrast, stimulation consistently increased ratings of physical pain over trials.

### Mood change and adverse effects

As shown in Table 2, Stimulation had no effect on mood change as measured by the VAMS (negative: F(1, 50) = 0.39, *p* = .54, ηₚ² = 0.01; positive: F(1, 50) = 0.71, *p* = .40, ηₚ² = 0.01) and did not interact with Stimulation site (negative: F(1, 50) = 0.73, *p* = .40, ηₚ² = 0.01; positive: F(1, 50) = 1.24, *p* = .27, ηₚ² = 0.02).

**Table 2.** Mood change and Adverse effects across stimulation sites and stimulation types. VAMS scores represent a change from pre to post levels of mood.

|  | rIFG |  | dmPFC |  |
| --- | --- | --- | --- | --- |
|  | Sham | Anodal | Sham | Anodal |
|  | M (sd) | M (sd) | M (sd) | M (sd) |
| VAMS-negative | -1.35 (5.79) | -0.01 (5.68) | -0.49 (3.97) | -0.71 (5.31) |
| VAMS-positive | -2.02 (3.41) | -2.19 (2.92) | -1.73 (2.14) | -0.47 (2.76) |
| Adverse Effects | 4.42 (2.94) | 4.54 (3.42) | 3.58 (3.37) | 3.62 (2.58) |

There was also no effect of Stimulation on adverse effects, F(1, 50) = 0.05, *p* = .83, ηₚ² < 0.001, and no interaction with Stimulation site, F(1, 50) = 0.01, *p* = .91, ηₚ² < 0.001.

## Discussion

Building on research showing overlapping neural processes associated with personal experiences of physical and social pain, it has been claimed that empathy for physical and social pain in others also relies on similar underlying neural processes (Iannetti et al., 2013). However, to date, causal evidence has been lacking. The aim of the current study was to provide causal, site-specific evidence for the roles of both the rIFG and the dmPFC in empathic judgements for social and physical pain in others using focal tDCS. Our results support a site-specific, dissociable role for the rIFG with excitation resulting in decreased ratings of social pain and increased ratings of physical pain.

The results of the current study are consistent with previous research implicating the rIFG in down-regulating the experience of social pain in others (He et al., 2018, 2020). Using both electrical and magnetic stimulation, He and colleagues (2018, 2020) presented images of social exclusion and then required participants to consider alternate explanations for the scenarios, a process labelled ‘reappraisal’. They showed that excitation of the broader rIFG region increased the ability to reappraise the socially painful scenarios using alternate, less painful explanations.

Our results, alongside those of He and colleagues (2018, 2020), support a causal role for the rIFG in emotional regulation that is specific to social scenarios. The broader rIFG region has been implicated in providing social meaning rather than considering physical features of an object or interaction (Martin et al., 2016; Tylén et al., 2016). Likewise, in primates, a synonymous brain region to the human rIFG was active exclusively during social interactions compared against physical interactions (Sliwa & Freiwald, 2017). The results of our study, therefore, add to a growing body of evidence highlighting the specialized role of the rIFG in processing and regulating socially relevant emotional experiences. This specificity underscores the importance of the rIFG as a neural substrate for navigating complex social interactions Our findings align with research indicating that processing social pain in others engages neural pathways distinct from those involved in perceiving physical pain, likely reflecting pathways associated with affective processing and regulation. For example, Woo and colleagues (2014) identified separable neural representations for pain and rejection and found activity in the broader right inferior frontal gyrus (rIFG) was negatively correlated with activity in the dorsal anterior cingulate cortex (dACC), a region consistently implicated in the experience of personal social pain. This relationship suggests that the rIFG may play a regulatory role in modulating the affective experience of social pain, potentially through top-down inhibitory control over the dACC. The rIFG has consistently been associated with response inhibition (Aron et al., 2014) and this extends to inhibiting or regulating emotions (Ochsner et al., 2012). Such findings highlight the possibility that rIFG activation is not only crucial for perceiving social interactions but also for regulating affective responses to social stimuli. Further research is needed to determine whether functional connectivity between the rIFG and the dACC underlies emotional regulation processes directed at social pain experienced by others. Investigating this interaction could provide valuable insights into the neural mechanisms that support empathy and social cognition, shedding light on how individuals navigate complex emotional landscapes in social contexts. Additionally, understanding these pathways may inform interventions aimed at improving emotional regulation in conditions characterized by heightened sensitivity to social pain, such as social anxiety or depression (Hudd & Moscovitch, 2020; Kupferberg & Hasler, 2023).

The rIFG is also considered a key hub of the hMNS (Betti & Aglioti, 2016; Gallese, 2007) with evidence for involvement in the processing of physical pain in both oneself and others (Baird et al., 2011). Neural activation in the rIFG has been shown to increase in response to observing others in pain (Budell et al., 2010, 2015). Crucially, after disruption of the rIFG using TMS, participants were slower to perceive pain in others (Li et al., 2021). To our knowledge, our study is the first to investigate whether excitation of the rIFG results in an increased perception of physical pain in others. Overall, our results are broadly consistent with previous evidence, with excitation of the rIFG increasing ratings of physical pain in others. However, we find no evidence that the increase in pain ratings is limited to painful images and instead show a general increase in the ratings of physical pain regardless of whether the image was depicting physical pain or not.

Although stimulation effects were greatest for the images depicting physical pain, there was no statistical difference in increased ratings when compared against the images depicting no physical pain. This suggests that stimulation to the rIFG increases the comprehension of pain in others regardless of whether they are experiencing pain or not. One possibility is that rIFG activation may signal potential threat as our no physical pain images depicted *potential* for pain, such as a near miss with a knife or sharp objects next to a bare foot. Previous research has found causal associations between the rIFG and threat detection (Clark et al., 2012), which has been explained as an increase in attention (Coffman et al., 2012) and perceptual sensitivity (Falcone et al., 2012), which suggests that rIFG activation may heighten awareness of environmental stimuli associated with potential harm. This enhanced attentional and perceptual processing could lead participants to interpret even innocuous images, such as a hand near scissors or thumb tacks, as threatening. Consequently, rIFG excitation may amplify the subjective experience of discomfort or anticipated pain, even in the absence of actual physical harm, aligning with its role in detecting and responding to potential threats.

The dmPFC is a region consistently associated with higher-order social cognition (Schurz et al., 2014), including empathy (Engen & Singer, 2013). It has been proposed as a potential target for neurostimulation treatments to improve empathic function in conditions such as frontotemporal dementia (FTD; Phillips et al., 2023). Previous brain stimulation research has focused on different aspects of self-other processing, demonstrating that excitation of the dmPFC can increase the salience of others, integration of social information, and emotion recognition (Ferrari, Lega, et al., 2016; Ferrari, Vecchi, et al., 2016; Gamond & Cattaneo, 2016; Martin, Dzafic, et al., 2017; Martin, Huang, et al., 2017, 2019). However, we found no effects of stimulation on empathic ratings of physical or social pain. The lack of an effect could be methodological, since the position of the dmPFC within the midline makes it harder to reach via electrical stimulation. This was confirmed by the theoretical current modelling with a reduced peak electrical field of 0.0946 at the dmPFC compared with 0.129 at the rIFG. Alternatively, the lack of an effect may be due to the nature of the empathy task used. The dmPFC has been associated with empathy but the majority of evidence points to a causal role in higher-order cognitive components of empathy. As the task used here focused on the affective component of pain in others (elicited by simple, static images), it could explain the lack of effects. Similarly, the dmPFC has been associated with the distinction and integration of self-other representations (Martin, Huang, et al., 2019; Wittmann et al., 2021) and may only be required for tasks that involve online processing of self-other information.

We anticipated a general habituation to both socially and physically painful images, expecting pain ratings to decline across the task. However, this pattern emerged only for social pain images; in contrast, ratings for physical pain images increased over the task. While not the primary focus of this study, this suggests that empathic responses to others in social pain may diminish over time. Neural habituation, a fundamental form of plasticity (Wilson & Linster, 2008), is adaptive for reallocating resources to novel stimuli. In the context of empathic responses, habituating to others’ pain may offer an evolutionary advantage when the pain cannot be alleviated. While neural habituation to pain has been demonstrated (Coll et al., 2016; Preis et al., 2015), behavioural habituation in the form of declining pain ratings has not been consistently observed (e.g., Lamm et al., 2007). To our knowledge, this study provides the first evidence of behavioural habituation differences in the empathic rating of social versus physical pain. Future research should investigate neural habituation differences for the observation of physical and social pain in others.

The exploratory analysis of stimulation response across trials revealed a differential effect of stimulation on social pain ratings over time. While excitatory stimulation to the rIFG consistently increased ratings of physical pain, it disproportionately reduced social pain ratings during earlier image presentations. One explanation involves differences between online and offline stimulation. During the first 20 minutes of the task, participants received online stimulation, which directly affects neuronal processes by depolarizing underlying neuronal populations. This facilitates neuronal firing when the area is engaged. Following cessation of active current, the offline period may reflect early neuroplastic changes (Liebetanz et al., 2002). Our findings tentatively evidence an online specific effect of stimulation to the rIFG for the reduced perception of social pain in others. This suggests that stimulation had no effects on the neuroplastic response at the rIFG and relied active, online stimulation to induce an effect. Although this requires further investigation and replication, the results have implications for the potential therapeutic use of f-tDCS in the regulation of social pain.

Alternatively, the rIFG may naturally habituate to repeated presentations of socially painful images. Neural habituation is an adaptive process whereby the brain reduces its response to repetitive or familiar stimuli, allowing cognitive and emotional resources to be allocated more efficiently to novel and salient events (Summerfield et al., 2008). In this context, habituation of the rIFG could lead to progressively diminished engagement with socially painful stimuli as the task continues, which, in turn, may explain the reduced efficacy of stimulation during later trials. This interpretation is consistent with evidence of neural habituation to painful images, particularly in the right insula and extending to the rIFG, as demonstrated by Preis et al. (2015). However, it remains unclear whether the habituation patterns for social and physical pain differ at the neural level. The specific role of the rIFG in processing social scenarios, such as reappraising social pain or providing social meaning, may explain why it habituates differently to social pain compared against physical pain. The top-down regulation of social pain by the rIFG may decline alongside the declining novelty of the stimuli, reflecting increased cognitive efficiency and reduced cognitive control requirements (Grill-Spector et al., 2006). Exploring these differences is essential to understanding the distinct neural circuits underlying empathy for social and physical pain. Such research could reveal whether the observed habituation is a general phenomenon of diminished affective engagement or a response specific to the cognitive and emotional demands of understanding social pain in others. Future studies using neuroimaging should investigate the time course of rIFG activity during tasks involving repeated exposure to social and physical pain stimuli. Examining connectivity changes between the rIFG, the insula, and other emotion-related regions, such as the amygdala and dorsal anterior cingulate cortex (dACC), could provide deeper insights into how habituation mechanisms differ across these two domains of empathy. These findings would not only elucidate the neural basis of habituation in social cognition but also inform interventions aimed at enhancing emotional regulation in disorders characterized by dysregulated responses to social or physical pain.

Although the use of real-life photos depicting pain and no-pain is more ecologically valid than the use of cartoon depictions or context-free facial expressions, future research could focus on increasing the ecological validity further. Future research should employ dynamic stimuli or depict more complex social interactions to assess whether findings from controlled lab-based environments are applicable to real-life scenarios that elicit empathic responses. Social neuroscience is increasingly moving towards a ‘second-person’ neuroscience (Redcay & Schilbach, 2019) which involves the investigation of real-time interaction at both the behavioural and neural level. For example, ‘second-person’ neuroscience methods have identified neural synchronisation as a key mechanism relevant for social cognition and interaction (Müller et al., 2021). Intriguingly, the rIFG has emerged as a region showing high levels of interpersonal synchrony (Jasmin et al., 2016; Shamay-Tsoory et al., 2024) pointing to a key role in social connectedness and interpersonal alignment. The results of the present study should motivate the adoption of such methods to further understand the role of the rIFG in empathic responses for social and physical pain in others, using ecologically valid, ‘second-person’ neuroscientific methods.

In conclusion, this study establishes a site-specific, causal, and dissociable role of the rIFG in empathic responses to social and physical pain in others. We show that rIFG excitation reduces ratings of social pain, whereas ratings of physical pain increase both when the image depicts pain and when there is potential for pain. In contrast, we find no evidence for a causal role of the dmPFC in empathy ratings for either social or physical pain. Exploratory analysis across trials revealed a disproportionately stronger stimulation effect on social pain ratings for images presented earlier in the sequence. These findings advance our understanding of the relationship between key social brain regions and empathy, challenging previous accounts of shared neural processes for empathy for social and physical pain. They also highlight exciting opportunities for future research employing more ecologically valid tasks that emphasize social interactions between two or more agents.

## Notes

### Competing Interest Statement

The authors have declared no competing interest.

